# Semantic computing for human phenotypes

**DOI:** 10.1101/2024.06.21.599369

**Authors:** Robert Gentleman, Rafael Goncalves, Vincent Carey

## Abstract

In many fields, research progress may be hindered by indefiniteness of language used to describe experimental conditions and outcomes. Harmonization of data resources generated by independent groups is important for integrative analysis. Adoption of formal ontologies and vocabularies for experiment annotation should help with harmonization tasks, but the use of ontologies also suffers from a lack of definiteness. In this study we explore how natural language characterization of human diseases coupled with ontologic mapping of study outcome terminology can be used to integrate information from multiple studies of genetic origins of disease risk. Open source tools and workflows are presented. This work exposes areas for improvement in tooling for data harmonization, which is a fundamental requirement for efficient research progress.

## Introduction

Larger and more complex biomedical data sets are becoming increasingly available and they provide substantial opportunities for integration across disparate data sources to enable the discovery of medically relevant relationships. Tools for representing and organizing biomedical entities and concepts are key components of any system that processes disease, medical or epidemiological data. Realizing the potential for discovery is hampered by the complexity of the data and the inherent challenge of working with phenotypes and experimental factors described primarily in natural language. To work with these textual descriptions there are a plethora of ontologies and vocabularies that are in use such as Unified Medical Language System (UMLS), (Bodenreider 2004), Medical Subject Headings (MeSH), Human Phenotype Ontology (HPO), (Gargano *et al*. 2024), and the Experimental Factor Ontology (EFO), (Malone *et al*. 2010). Finding and grouping or comparing phenotypes of research interest in large data resources such as NHANES, (CDC 2024), or biobanks, such as the UKBB, (Sudlow *et al*. 2015), can be aided by effective use of ontologies and tools for natural language processing.

We explore some of these challenges in the context of the GWAS Catalog, (Sollis *et al*. 2023), which reports on a set of published curated GWAS association studies where the phenotypic traits reported in published articles are mapped to terms in the EFO. A GWAS reports on assocations between single nucleotide polymorphisms (SNPs), which are variants in the human genome, and specific traits or phenotypes. The GWAS Catalog provides an integrated database of a large set of published GWAS studies curated from the scientific literature and collated, together with a large amount of metadata. Users can freely download and search the data for different purposes. While the data are freely available, access to them through easy to use interfaces designed for non-experts appears to be mainly through search facilities on the GWAS Catalog website. Our interest is in integrating this resouce with search facilities in the R programming language in order to provide programmatic access to the data and to enable the sorts of complex searches that would be of interest in epidemiological and other fields.

To facilitate searching for traits we use a search strategy that combines the use of a dictionary (corpus based) search combined with an ontology based search. We create a dictionary search capability that is built on the textual descriptions from both the GWAS Catalog and EFO which we combine with a set of tools designed to make use of the relationships and concepts in the ontology. We demonstrate how these tools, when used together, enable programmatic search of the GWAS Catalog. The approach has some similarity to that of phenotagger, (Luo *et al*. 2021), a method that combines both dictionary and machine learning-based methods to identify (HPO) concepts in unstructured biomedical text.

In particular we examine the use of corpus search tools, as exemplified in the `corpustools` package, (Atteveldt 2023), to help search the set of traits that are reported on by the GWAS Catalog. Since these investigator defined traits have been hand curated by the EBI and mapped to terms in the EFO we also examine how that mapping can be used to aid the analysis and interpretation of the data. Our software is contained in an R package, `gwasCatSearch` (Gentleman, Carey and Gonçalves 2024), which is open source and freely available.

## Materials and Methods

### Data

Since 2010, delivery and development of the GWAS Catalog has been a collaborative project between the EMBL-EBI and NHGRI. Full details of the data and the date it was obtained are given in Supplementary Data. GWAS Catalog data is currently mapped to Genome Assembly GRCh38.p14 and dbSNP Build 156. The curation process used is described at https://www.ebi.ac.uk/gwas/docs/methods/curation. As of Feb 6, 2024, the GWAS Catalog contains 6679 publications and 571148 associations. The results reported in a publication are further divided into *studies* with unique accession identifiers. Each publication can report one or more studies. The online text on the curation process includes the remark: *Each study is also assigned a trait that best represents the phenotype under investigation. When multiple traits are analysed in the same study either multiple entries are created, or individual SNPs are annotated with their specific traits*.. This indicates that in some situations individual SNPs will be annotated with a trait and unique study accession number, while in other cases collections of SNPs will be associated with a single trait and accession number.

We obtain data from the GWAS Catalog and transform it into a set of tables using tools in the GWASCata-logSearchDB (Gonçalves and Payne). Our processing produces a number of tables that can be accessed via a SQL database; we use SQLite (Hipp 2020) following the example of the INCAtools project (Matentzoglu *et al*. 2022). These tables include data from the EBI, several tables organized around the Experimental Factor Ontology (EFO) and the Uber-anatomy ontology (UBERON) (Mungall *et al*. 2012).

### Ontologies

An ontology is a set of concepts and categories that describe a subject area or domain of knowledge that expresses their properties and the relations between them. Ontologies form one of the bases for knowledge engineering and representation and are important in understanding and referencing biomedical entities. One of the challenges that needs to be addressed is the fact that the same concept might usefully appear in different ontologies and users will need tools to help identify similar concepts across ontologies. A second challenge is that to be useful the ontologies must be accessible and typically that means using the web or a database implementation. There are two constructs that help with this, the compact uniform resource identifiers (CURIEs), and the internationalized resource identifiers (IRIs). A CURIE has three parts: a prefix; a delimiter; and a unique local (to the ontology described by the prefix) identifier from the given nomenclature. The CURIE EFO:0005305 which represents the concept *atrioventricular node disease* has a prefix of EFO, a delimiter of a colon (:) and a unique local identifier of 0005305. The corresponding IRI for that term is http://www.ebi.ac.uk/efo/EFO_0005305, which gives a web location that can be examined by a user. Note that the delimiter in the IRI is an underscore, rather than a colon. We will use both in our discussion. The terms in the EFO ontology are related to each other via semantic relationships, which can be viewed as a directed acyclic graph. By construction there is a single root node. More specific terms, such as the relationship between cancer and pancreatic cancer are encoded with pancreatic cancer being the **child** of cancer, while cancer is the **parent** of pancreatic cancer. The descriptions can become more refined, and we define the **descendants** of a node to be the set of child nodes, where one recurses down to the point where a node has no children. Such a node is referred to as a **leaf**. Studies that map to leaf nodes in the ontology are describing quite specific concepts, while studies that map to non-leaf nodes address concepts that are more general. Similarly we define **ancestors** to be those nodes obtained by recursing the parent operation applied recursively up to the root node of the ontology.

Our discussion of ontologies focuses on term collections that employ `is-a`, `part-of`, and `has-disease-location` to relate terms. Given a term in an ontology, the operations `parent-of`, `children-of`, `descendants-of` have natural interpretations. The root node of the ontology is the parent of itself.

Our software contains tools to obtain the children, descendants, parents, and ancestors as well as to perform other operations using the ontology. Given the structure of the ontology we can introduce the concept of an *inherited* mapping. A mapping at a node E is inherited if it was a direct mapping to some descendant of E. For example the CURIE for pancreatic carcinoma is EFO:0002618, which has 44 direct mappings and 1 inherited mapping. (The inherited mapping is for a study (Pistoni *et al*. 2021) that mapped to pancreatic ductal carcinoma.) The structure of the ontology provides substantial benefits with regard to the organization of phenotypes, as, for example, all diseases should be organized under the node labeled *Disease* which has CURIE EFO:0000408. Thus, by examining the direct and inherited mappings at the EFO:0000408 node we can identify all GWAS Catalog traits that have been mapped to a human disease. Now, that will not be quite as useful as it seems due to issues we will explore due to the inherent complexity of phenotypes.

Information on anatomic location of diseases was extracted directly from EFO statements of the form: X has_disease_location Y, where Y is typically an UBERON term representing an anatomical location. If a term does not have an explicitly stated location relationship, we determine if it has an inferred location. This is done by recursively checking if an ancestor in the ontology hierarchy has a location. The first location found is used, and if none is found then it is marked as missing. For example, *bronchitis* (EFO:0009661) does not have an explicitly stated location in EFO, but it inherits one from its immediate parent (*bronchial disease*), which has location *bronchus*, (UBERON:0002185). Of the 39617 EFO terms 6346 terms have a disease location.

Our capacity to identify anatomic locations of anatomically localized diseases is limited to some extent by logical practices of ontology designers. In some cases, the disease location is provided as a statement of the form: kidney disease has_disease_location (kidney or part_of kidney). Logically, this does not entail that kidney disease has_disease_location kidney. The statement does not specify the location. Thus we miss the locations of diseases that are defined this way in EFO. Sometimes the ontology developers did add an extra statement that would unambiguously give us a location. For example, for bronchial disease, which is defined as a disease that has_disease_location (bronchus or part_of bronchus), there is an additional statement that bronchial disease has_disease_location bronchus. If this pattern were consistent, we would get more comprehensive coverage of locations. Inference of disease location from disjunctions, in which (A `or part-of` A) can be reduced to A as a location, could be supported with a Web Ontology Language inference engine, but we have not undertaken this.

### Aligning EFO and the GWAS Catalog

To facilitate searching we create a dataframe based on the GWAS Catalog studies in which there is one row for each unique combination of GWAS Catalog study ID and EFO term. This GWAS Catalog dataframe has columns labeled `STUDY.ACCESSION`: the GWAS Catalog study identifier; DISEASE.TRAIT: the GWAS Catalog text description of the trait; `MAPPED_TRAIT`: the textual description of the trait from EFO that this study was *directly* mapped to; `MAPPED_TRAIT_URI`: the URI for the mapped trait; `MAPPED_TRAIT_CURIE`: the CURIE for the mapped trait. There are 89958 studies reported. Each study can be mapped to one or more EFO terms. We counted the number of EFO terms each study was mapped to and report those in Table 1. The counts range from 1 to 58 with the study GCST001762 having the most and the vast majority of GWAS Catalog studies are mapped to only one EFO term.

**Table 1:**
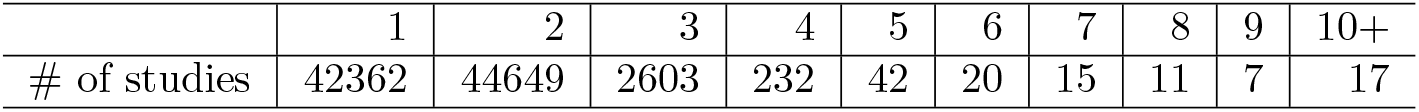
GWAS Catalog Studies by number of EFO terms per Study.

|  | 1 | 2 | 3 | 4 | 5 | 6 | 7 | 8 | 9 | 10+ |
| --- | --- | --- | --- | --- | --- | --- | --- | --- | --- | --- |
| # of studies | 42362 | 44649 | 2603 | 232 | 42 | 20 | 15 | 11 | 7 | 17 |

We also provide a dataframe representing the EFO terms, with one row for each term in the EFO ontology and columns labeled `Subject`: the corresponding EFO CURIE; `Object`: the textual description of the trait; `IRI`: the IRI for that term; `DiseaseLocation`: the UBERON CURIE for the location of the `trait` if there is one; `Direct`: the number of GWAS Catalog studies that were mapped directly to that trait; `Inherited`: the number of traits that were mapped to terms that are more specific (according to the ontology) than the `Object`.

The contents of the two dataframes are summarized in Tables 2 and 3. For the EFO data there are two columns that were not included in the summary table, `Direct` which contains the count of GWAS studies that were directly mapped to that trait and `Inherited` which is the count of GWAS studies that are inherited from children of the EFO term.

**Table 2:** Overview of GWAS Catalog Data.

| Field | Unique values |
| --- | --- |
| STUDY.ACCESSION | 89958 |
| DISEASE.TRAIT | 67152 |
| MAPPED_TRAIT | 18086 |
| MAPPED_TRAIT_CURIE | 16066 |
| MAPPED_TRAIT_URI | 16063 |
| MappingScore | 401 |
| Source | 2 |
| Tags | 2 |

**Table 3:** Overview of EFO Data.

| Field | Unique values |
| --- | --- |
| Subject: EFO CURIE | 39617 |
| Object: Trait Name | 39612 |
| IRI: IRI | 39617 |
| DiseaseLocation: UBERON CURIE | 171 |

In Table 4 we list the EFO terms that have the highest frequency of GWAS studies mapped to them. And considering the results in Table 4 in combination with those reported in Table 2 we can infer that for many of the GWAS Catalog studies there is relatively little information contained in the EFO mappings as the EFO terms listed in Table 4 are quite generic.

**Table 4:**
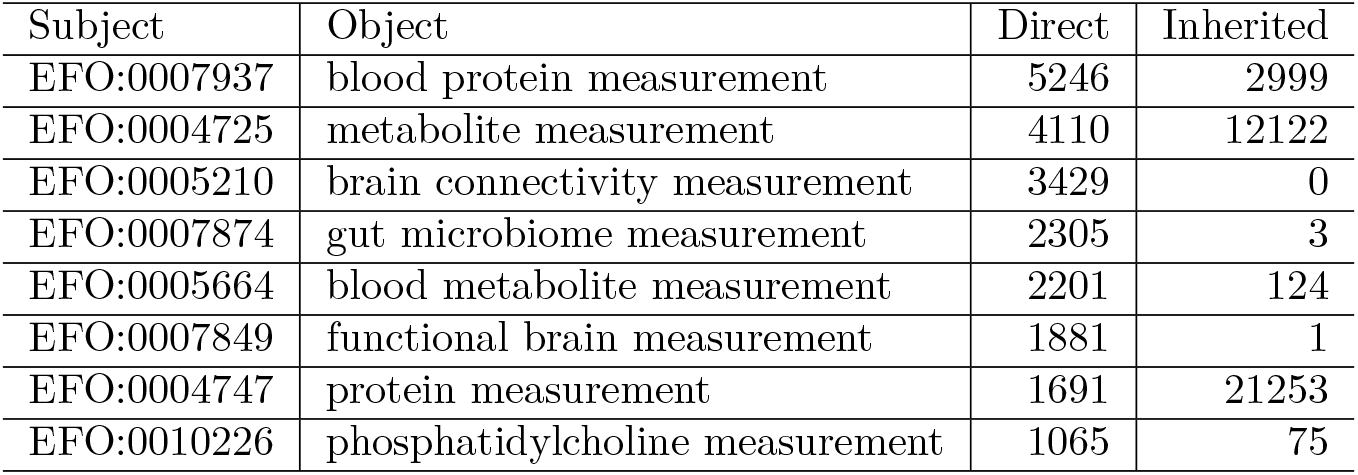
The top 8 records in EFO based dataframe, ordered by frequency of direct annotation to GWAS.

The GWAS Catalog metadata table contains one entry per STUDY.ACCESSION number and other variables such as the PUBMEDID for the associated publication, more complete information on these tables is in the Appendix. We now examine the relationship between published papers and the GWAS Catalog studies. A single paper can give rise to more than one GWAS Catalog study. We find that there are 4276 papers for which a single GWAS Catalog study is identified. On the other end, we find that there are 68 publications which have more than 100 GWAS studies associated with them. We examined the paper associated with PMID 34662886 (Backman *et al*. 2021) which has 7972 GWAS Catalog studies associated with it. In the Backman *et al* paper the authors report “We identified 12 million coding variants, including around 1 million loss-of-function and around 1.8 million deleterious missense variants. When these were tested for association with 3,994 health-related traits, we found 564 genes with trait associations at *p* ≤ 2.18 *×* 10 − 11. So it does seem likely that these 7972 studies do represent distinct traits.

### Building search corpora

To create a searchable EFO-based corpus we create synthetic documents that consist of three components: first the EFO term’s text description; second the EFO provided synonyms; and third the set of GWAS Catalog study names, for any study that was directly mapped to that EFO term. Our rationale for the third component is that the EBI curators assigned that trait description to the specific EFO term, so it is in their opinion a synonym. We show the EFO provided synonyms for kidney disease, EFO:0003086 in Table 5.

**Table 5:** Synonyms for EFO:0003086, kidney disease.

|  |  |
| --- | --- |
| renal disorder | disease of kidney |
| kidney disorder | renal system disease |
| kidney disease or disorder | renal disease |
| kidney disease | nephropathy |
| disorder of kidney | kidney diseases |
| disease or disorder of kidney | Kidney Disorder |

For clarity, we create one *document* for each EFO term. The document has three components, the term itself we call the *object*, second a set of zero or more *synonyms* provided by EFO and third a set of zero or more *matches* which are GWAS study traits that have been mapped directly to the EFO term.

Searching for relevant GWAS studies using this corpus one first obtains all EFO terms where the tokenized search string matches at least one of the *object, synonyms* or *matches* for that term. The desired GWAS studies are those that have either matched directly to that EFO term, or are inherited by that term. Suppose our search string had been *chronic kidney disease*, then we would have found the term EFO:0003086, which has as its text description *chronic kidney disease*. There were 16 GWAS Catalog studies that were mapped **directly** to this EFO term and 593 studies that were inherited, i.e. mapped to more specific EFO terms than EFO:0003086, which could be subtypes.

In Figure 1 we render a portion of the EFO ontology that is anchored at the node labeled urinary system disease, EFO:0009690, and shows all of the descendants of that term.

**Figure 1.**
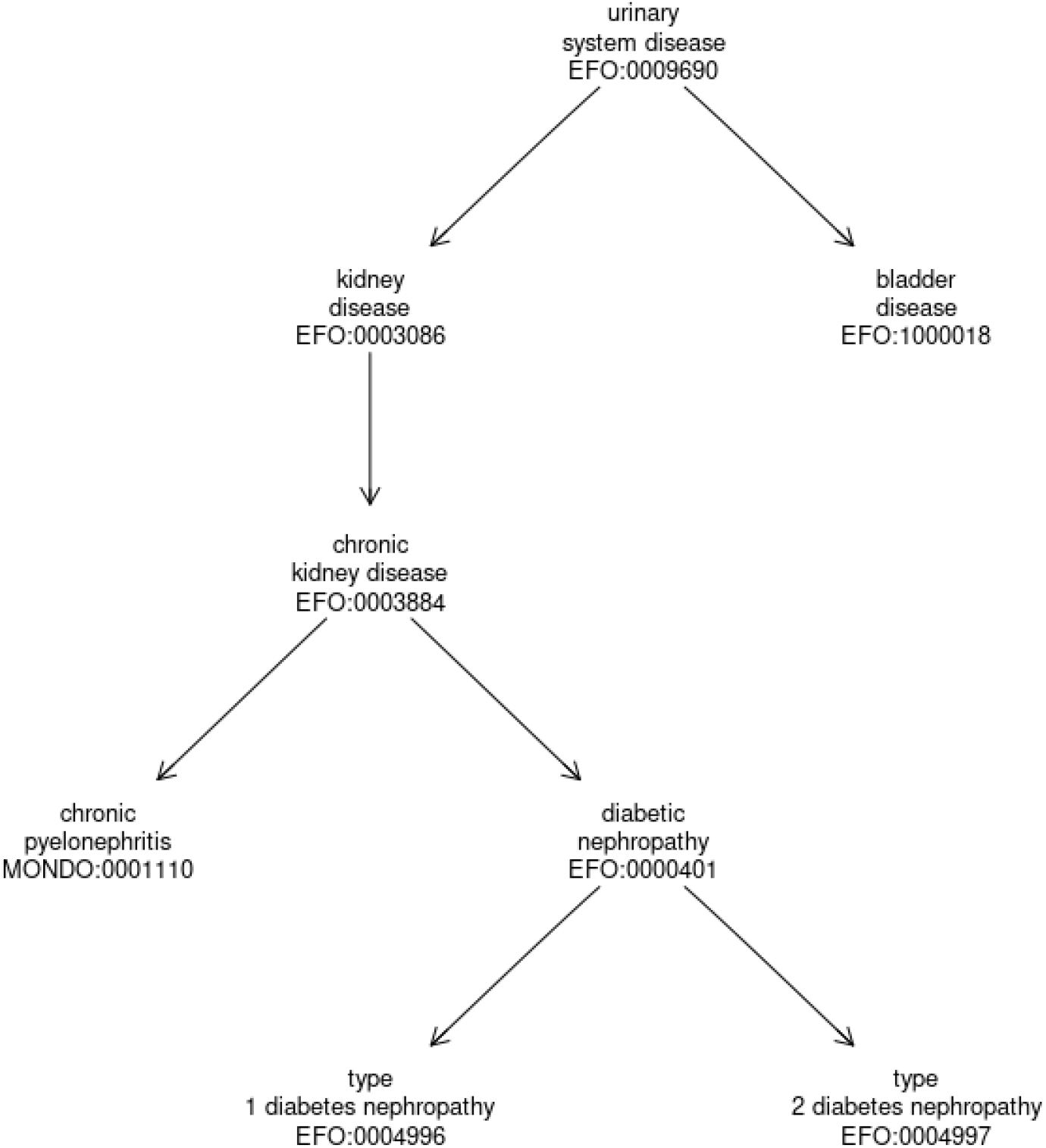
Rendering of the EFO Ontology for urinary system disease and descendants.

### A search corpus based on GWAS Traits

To create this searchable corpus we made documents, one for each GWAS study, by assembling the GWAS trait (DISEASE.TRAIT), the EFO mapped trait (Trait) and all EFO provided synonyms (Synonyms). In this case the ontology is being used to provide a standardized trait name and its synonyms. Relationships within the ontology are not used. The corpus can be searched and studies that match the search term retrieved. In Table 6 we show the metadata associated with the study GCST90083943 as well as the synonyms for EFO:1001455. The entry in our corpus would consist of the text strings associated with the entries DISEASE.TRAIT, MAPPED.TRAIT and EFOSynonyms in that table.

**Table 6:** Some attributes of Study GCST90083943 in the GWAS Catalog.

|  |  |
| --- | --- |
| STUDY.ACCESSION | GCST90083943 |
| DISEASE.TRAIT | ICD10 H93: Other disorders of ear, not elsewhere classified (Gene-based burden) |
| MAPPED_TRAIT | auditory system disease |
| MAPPED_TRAIT_CURIE | EFO:1001455 |
| MAPPED_TRAIT_URI | http://www.ebi.ac.uk/efo/EFO_1001455 |
| MappingScore | NA |
| Source | GWASCatalog |
| Tags | None |
| EFOSynonyms | disorder of auditory system disease or disorder of auditory system; disease of auditory system auditory system disorder; auditory system disease or disorder auditory system disease; auditory disease otological disease; otologic disease ear disorder NOS; ear disorder ear disease; ear and mastoid disease disorder of ear; Unspecified disorder of ear Destruction - ear drum |

The same set of textual descriptions are used for constructing both this corpus and the EFO corpus described above. These are the original set of traits as defined by the authors of the studies and the traits and synonyms for all EFO terms that were mapped to. So a search based on either one of these should yield the same set of EFO terms and GWAS Catalog studies. We find that it is useful to have both.

As shown in Table 7 it is also possible to identify which of the three parts of the *document* described above have matches. There were 552 unique documents that were identified. We can see that there were 275 cases where a match was to the DISEASE.TRAIT, 4426 matches to synonyms provided by the ontologies, and 699 matches to the text description used by the ontology for the trait.

**Table 7:** Which part of the GWAS *document* matched the search token leuk*.

| Term | Count |
| --- | --- |
| DISEASE.TRAIT | 275 |
| Synonyms | 4426 |
| Trait | 699 |

### Data Availability

All data used in this paper are publicly available, as are all softwore tools. Specific URLs and on-line resources are indicated at their point of use in the article.

## Results

### Facilitated search for diseases and traits

There are a number analytic tasks that these GWAS Catalog-based tools can aid in. A basic example is retrieval of associations reported on all diseases of a fairly broad class, say disorders of the kidney, or infectious disease. Another example is enumerating diseases and genetic associations that relate to some organ, system or part of the body. Having traits mapped to an ontology, such as EFO, should help with these tasks. We will examine how these tasks can be supported using the EFO ontology together with the search corpora we describe above. This process integrates a dictionary search with the structured information in an ontology.

We first consider the question of whether the GWAS traits are mapped to ontology terms that represent diseases. The EFO CURIE for *Disease* is EFO:0000408 and all diseases should be children of that node. We find that 93 GWAS studies were mapped directly to this node while 13335 were inherited. Since there are 89958 different GWAS studies, we have the surprising finding that less than 20 per cent of studies in the GWAS catalog present associations that were mapped to this part of the EFO.

The use of string matching for retrieval of genetic associations of potential interest can have unexpected consequences. We searched each assigned disease trait for any occurrence of the string *chronic kidney disease*, ignoring capitalization, and found 10764 matches which correspond to 37 different PUBMEDIDs. These studies are mapped to 8340 different EFO terms, with 823 mapping to EFO:0004725, which is *metabolite measurement*. We will return to this specific example in our discussion.

### A worked example: genetic associations with “infectious disease”

We will look for GWAS studies associated with *infectious disease*. One reason for this choice is that it is quite a challenging problem if one attempts it without using an ontology. The main reason for that is that there are many infectious diseases and most of them do not have either *infectious* or *disease* in their name. So without an ontology to guide us, we would need to have some dictionary of infectious diseases to start with, and for that there are several candidates, but we were unable to identify a definitive authoratative source.

#### Strategy One: Use the EFO ontology structure

Here we describe how to use these tools. We start with a general type of disease, say infectious disease, and then try to find all GWAS studies that report on that class of diseases. First we identify the node in the EFO ontology that corresponds to *infectious disease*, which has CURIE EFO:0005741. All infectious diseases should be annotated in the EFO as descendants of that EFO term. Currently there are 670 descendants and a total of 968 GWAS studies annotated at EFO:0005741 or one of its descendants. We find that 17 were directly mapped to EFO:0005741 while 951 where inherited. From Table 8 we see that there were 503 EFO terms that had no studies mapping to them. We note that the EFO term with over 200 studies mapped to is is COVID-19,

**Table 8:**
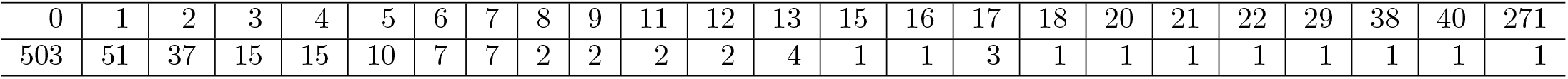
Number of studies mapping to descendants of EFO:0005741 (infectious disease)

#### Strategy Two: Search based on the GWAS Catalog corpus

In this case we will search the corpus based on the GWAS Catalog. One might want to experiment with different queries to try to obtain a good trade off between sensitivity and specificity. We found that using *infectious disease* as our query missed a number of GWAS studies that had *infection* in them and opted to use only the term *infect\**. If a list of all infectious diseases was available a different strategy would be to search for each disease in that list and then merge the results. Recall that when we constructed the GWAS Catalog corpus we used the text descriptions of the GWAS trait as well as the EFO trait name and synonyms for the EFO term(s) this study was directly mapped to.

We found 1031 GWAS Catalog studies and in Table 9 we report where in the *document* matches have occurred, when a match was in more than one part of the *document* then both are reported, we see that the majority of matches are to synonyms.

**Table 9:** Which part of the GWAS Catalog document matched our search term.

| Term | Count |
| --- | --- |
| disease.trait | 67 |
| disease.trait, synonyms | 56 |
| disease.trait, synonyms, trait | 129 |
| disease.trait, trait | 20 |
| synonyms | 566 |
| synonyms, trait | 169 |
| trait | 24 |

The second approach found 235 studies that were not found by our first approach, while the first approach found 172 studies that were not found by the second. One of the studies found by second approach that was not found using the first was GCST000346 which has the disease trait Cystic fibrosis severity. And the associated EFO term is MONDO:0009061 and while it is not an infectious disease one of its synonyms is “pseudomonas aeruginosa, susceptibility to chronic infection by, in cystic fibrosis”, which explains why it was found using our second method. Examining the list of those found by our first method but not by our second we see that these are mainly infectious diseases where the token *infect* is not found in the name of the disease or any of its synonyms.

#### Strategy three: Use the EFO corpus to get specific disease names

Above we first started by searching for *infectious disease* in the EFO ontology and then found all GWAS studies that mapped directly to either that node, or one of its descendants. We contrasted that with the strategy of searching the corpus based on the GWAS Catalog studies for the terms infectious and disease. A third approach is to use the EFO corpus to provide specific disease names, from the term labels for those EFO terms listed under the infectious disease node, EFO:0005741. If we simply extract all the EFO term labels for the descendants of EFO:0005741 and search the corpus with those we find 1170 studies, which is an increase over using the first strategy that relied on the direct and inherited studes reported for EFO:0005741. We could enhance thid search further in a number of ways such as using all synonyms, or breaking down terms into individual words and searching. However these processes can become unwieldy.

### Searching for GWAS studies by Disease Location

As a final example, we describe how one can make use of the disease location information provided by EFO. First, we must identify the UBERON CURIE that corresponds to the location of interest. The CURIE for lung in the UBERON ontology is UBERON:0002048 and we will use that to find EFO terms that specify UBERON:0002048 as the disease location. We identify 104 terms as having lung as their location. In order to provide some sense of the terms that were found we create a word cloud using functionality in the `corpusTools` package and visualize this in Figure 2.

**Figure 2.**
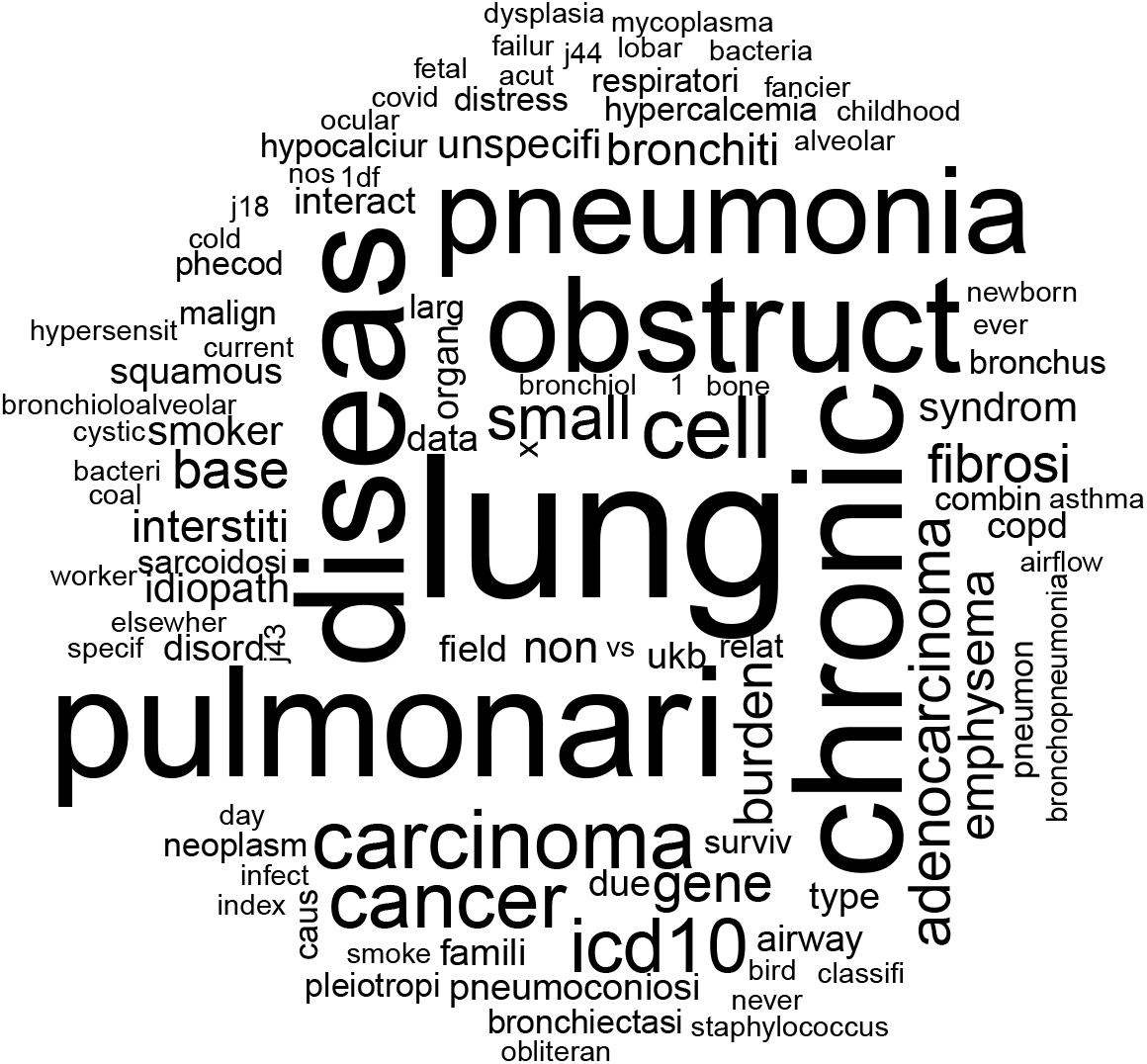
Word Cloud based word frequency in the EFO labels of those EFO terms that have disease location UBERON:0002048. The data were subset to only include tokens that have at least 5 occurrences.

Once we have a set of GWAS studies that we are interested in we can access the metadata for the implicated variants such as likely gene, the genome coordinates, the p-value and effect size to name a few of the attributes that are provided. These data are large and we use a database as the main tool for storing and accessing the data. The data can be read into standard Bioconductor objects, such as those for storing and manipulating genetic coordinates and users can easily produce Manhattan plots such as Figure 3, or identify genes, where for example we report the ten most frequently reported genes for this collection of studies in Table 10. Notice genes such as the cholinergic receptors CHRNA3 and CHRNA5 with strong associations for risk of lung cancer.

**Table 10:**
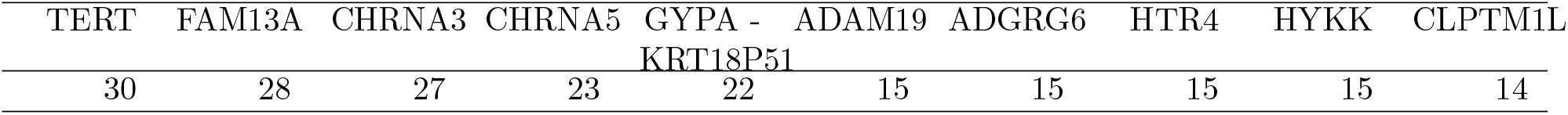
Most frequent mapped genes in GWAS lung studies.

**Figure 3.**
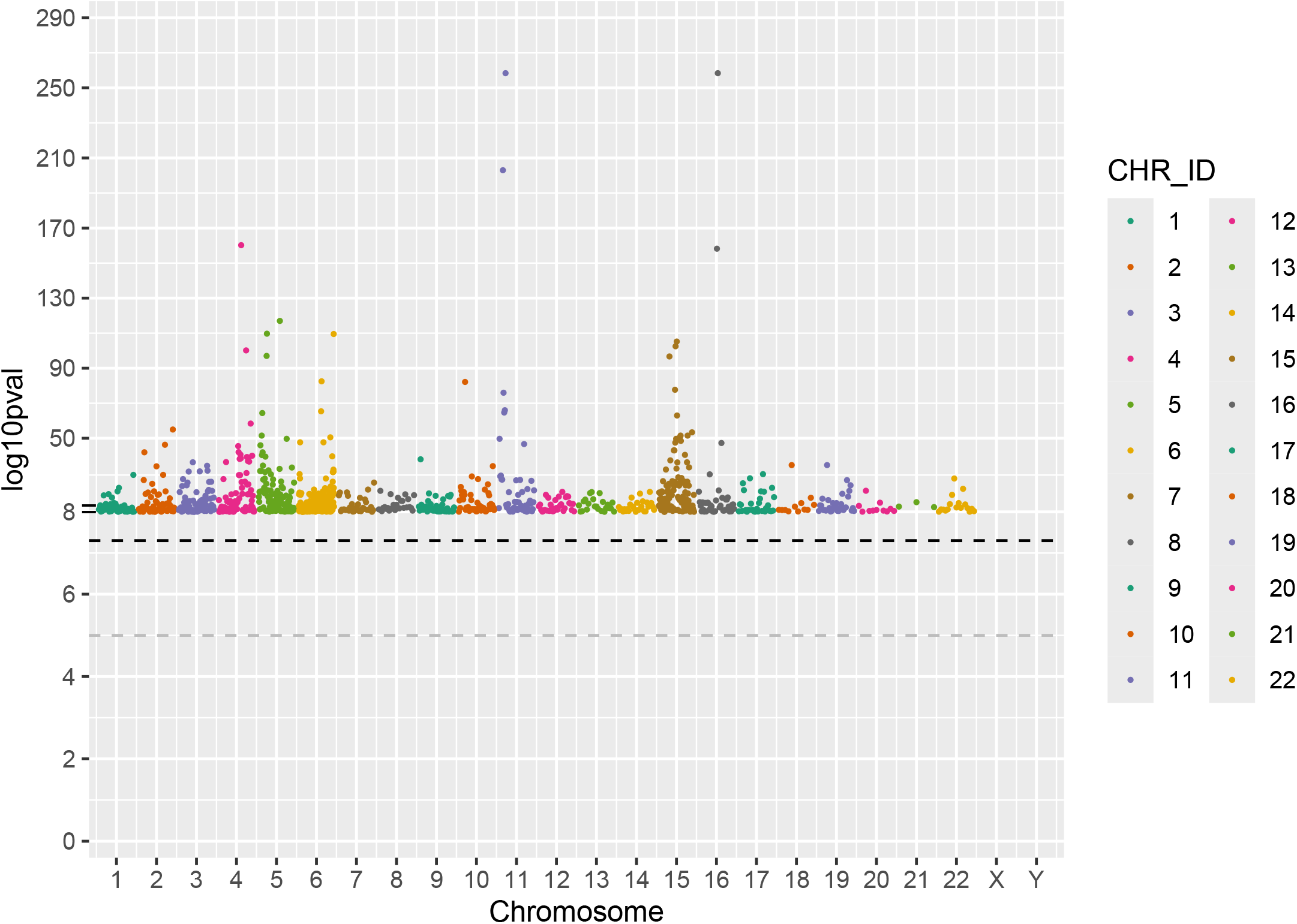
Manhattan Plot for GWAS variants associated with studies where we are able to identify the disease location Lung

### Consequences of imprecise annotation of study outcomes

While the GWAS Catalog has provided mappings to EFO there remain some substantial challenges for those attempting to use it to find and organize their searches. As we see in Table 4 a large number of GWAS studies are mapped to EFO terms where the EFO label is not particularly descriptive so that for these studies EFO labeling has contributed very little information. It is possible that the GWAS Catalog traits will contain more specific details, however, accessing that information will be reliant on tools for searching plain text.

The disease location data are useful, but the mapping is incomplete, since the predicates and search strategies that will need to be employed to access it for these sorts of analysis are complex. Expertise in the semantics of ontological concept representation and familiarity with the specific ontologies being used will be required. These are usually challenging for investigators in disease genetics.

We also found in the examples above that the study of the traits associated with a specific disease requires a fairly comprehensive search across all components of an ontology. For example, previously we discussed the term EFO:0003086 that has the text description: “chronic kidney diseases”. We found 16 GWAS Catalog studies that were mapped **directly** and 593 inherited studies for that EFO term. But these are not all the GWAS Catalog studies that have the string *chronic kidney disease* in their trait description. When we search the corpus based on the GWAS Catalog we find 10809 different studies that have a hit.

A little more exploring suggests that many of the terms are actually mapped to the descendants of the *measurement*, EFO:0001444, node. Specifically we find that there are 1472 EFO terms that were identified by our search which are annotated as measurements, a number far larger than those that align to disease. Thus, relying on mappings to the disease hieriarchy alone will miss many traits relevant to chronic kidney disease.

When we look at the PUBMED IDs associated with these traits we see that there are 3 papers that report more than 1000 traits. Given the PubMed IDs it is easy to find the papers. The paper associated with PMID 35870639 which is associated with more than 6,700 studies is titled: Identification of 969 protein quantitative trait loci in an African American population with kidney disease attributed to hypertension (Surapaneni *et al*. 2022).

Another potential issue that will affect the completeness and accuracy of our results is that the EFO provided synonyms might be incomplete or inaccurate. For rheumatoid arthritis the reported synonyms include *proliferative arthritis* and *Rheumatic gout*. Proliferative arthritis refers to a condition characterized by the excessive growth of synovial tissue in the joints, leading to inflammation and joint damage while rheumatoid arthritis is an autoimmune disease where the immune system mistakenly attacks the joints, causing inflammation, pain, and joint damage. And *Rheumatic gout* is a somewhat archaic term for gout. Including these as synonyms will result in linking traits that should not be linked.

## Discussion

The results of our exploration are encouraging but also exhibit the challenges that remain. The mapping of phenotypes to an ontology is extremely helpful in providing a more specific vocabulary that can be used to harmonize across studies. The terms or nodes in the ontology provide a means of identifying synonyms that can be incorporated into searches. The ontologies themselves are often undergoing active research and development. That typically means they will be updated and in some cases there are ways for users to provide feedback about issues and to give suggestions. As we have noted they can include additional information, such as disease location, that can be used in any analyses.

Although there are certain advantages in applying ontology labels to human phenotypes, the effectiveness of correlating them directly with specific phenotypes is debatable. Challenges arise when trying to accommodate concepts like the age of initial disease diagnosis, or the quantification of a protein level shift in a disease compared to a healthy state. There is a tension between accurately depicting the entity, such as age in the first instance and protein measurement in the second, and precisely pinpointing the root disease. Perhaps an approach where the phenotypes of interest are tagged, or labled as pertaining to certain concepts, measurements or diseases would facilitate broader and more appropriate usage.

There are no clear mechanisms to describe the curation processes used. And of course these may change over time, or be different for different curators adding to the complexity of relying on this strategy for aligning phenotypes with ontological representations of knowledge. We have shown that in the case of the GWAS Catalog and curation to the EFO curators chose to map phenotypes to entries in the *measurement* portion of the EFO ontology rather than the disease hierarchy. This is perfectly valid but as our use case relies on identifying diseases this decision leaves us relying on syntax and string matching. We anticipate that further refinements, such as making use of other natural language processing tools such as name-entity-recognition (NER), will be useful in further identifying phenotypes that reference specific diseases, genes, drugs.

We believe that it is clear that use of ontologies and similar tools for organizing and analyzing natural language (e.g. artificial intelligence tools for natural language processing, and name entity recognition tools) will play an important role in helping solve some of the existing problems. We also believe that the challenges are complex and require cross-disciplinary efforts with somewhat well stated analytic objectives. Our ability to integrate and synthesize data across studies remains hampered by other issues as well. In most situations we would like to be able to address whether or not studies were based on reasonably similar cohorts of individuals, and in some cases whether or not the inclusion and exclusion criteria were similar. These remain challenges across much of medical and epidemiological data reporting.

