## Supplement for "Semantic computing for human phenotypes"

### gwasCat Supplement

2024-04-27

#### The paper

Our paper was written using bookdown Xie (2024) and the markup is contained in the *gwasCatSearch@ package available on GitHub, https://github.com/ccb-hms/gwasCatSearch. Within that repository, in the vignettes folder are all the documents needed to reproduce this paper. All details are given explicitly in the file GWAScatPaper.Rmd and they can be modified and extended if the reader chooses to explore our analysis.

#### GWASCatalogSearchDB

This paper relies on a large amount of work to organize the GWAS catalog data resources in a manner that is suitable for the analyses we describe. The explicit details of that effort are available from the corresponding GitHub repository, https://github.com/ccb-hms/GWASCatalogSearchDB. We summarize the process here for interested readers.

We programmatically obtained two tables from GWAS Catalog's Downloads webpage https://www.ebi.ac.uk/gwas/docs/file-downloads: 'All studies v1.0.2' and 'All associations v1.0.2', and store them without modification as tables `gwascatalog_metadata` and `gwascatalog_associations`, respectively. The first table contains details about GWAS studies registered in the GWAS Catalog, while the second contains details about the SNP-trait associations extracted from those studies.

In the `gwascatalog_metadata` table, there are sometimes multiple ontology mappings for a single study, which are represented as a comma-separated list of CURIEs in each study's row. This representation makes search over such values challenging. So, we extract all ontology mappings and create another table, `gwascatalog_mappings`, where each ontology mapping is represented in its own row. The rationale behind this new table is that it can be extended to include additional mappings from different sources which allows users to select their preferred mapping source(s) to use for search.

From the `gwascatalog_metadata` we extract the PubMed ID associated with each study, and we use the `metapub` (https://pypi.org/project/metapub) Python package to extract publication details such as titles, abstracts, and journal names, which we then store in the table `gwascatalog_references`. These data could easily be embedded into a vector database that could then facilitate use in large language model (LLM) searches such as the retrieval augmented generation (RAG) approaches that are now becoming commonplace.

For the experimental factor ontology (EFO) our database contains a table named `efo_labels` with details about all terms in EFO, such as their labels, identifiers, and disease locations associated with them (if available). In this table we also include the count of how many GWAS studies are directly mapped to each ontology term, and of how many studies are indirectly mapped to each term, i.e., they are mapped to a more specific term in the hierarchy.

Our database includes a tabular representation of an ontology's class hierarchy before and after reasoning, in the tables `efo_edges` and `efo_entailed_edges`, respectively. So for a particular term, we can obtain its direct, asserted parents using `efo_edges`, and we can obtain its indirect, entailed parents (i.e., ancestors) using `efo_entailed_edges`. The table `efo_synonyms` contains the potentially multiple synonyms of each EFO term. We construct our tables by extracting the minimum required details from a SemanticSQL representation of EFO.

Our pipeline allows the inclusion of any additional user-specified ontologies. In our case, we include the Uberon ontology so that we can obtain more details about the anatomical structures associated with (EFO) diseases. The tables included in the database for such extra ontologies are the same as for the primary annotation ontology, except that the labels table does not include disease location details or mapping counts.

Finally we include a table in our database (version_info) specifying the versions of the ontologies used in the pipeline, and the dates and times when the metadata tables were downloaded from the GWAS Catalog.

A schema of the resulting tables is given in Figure 1.

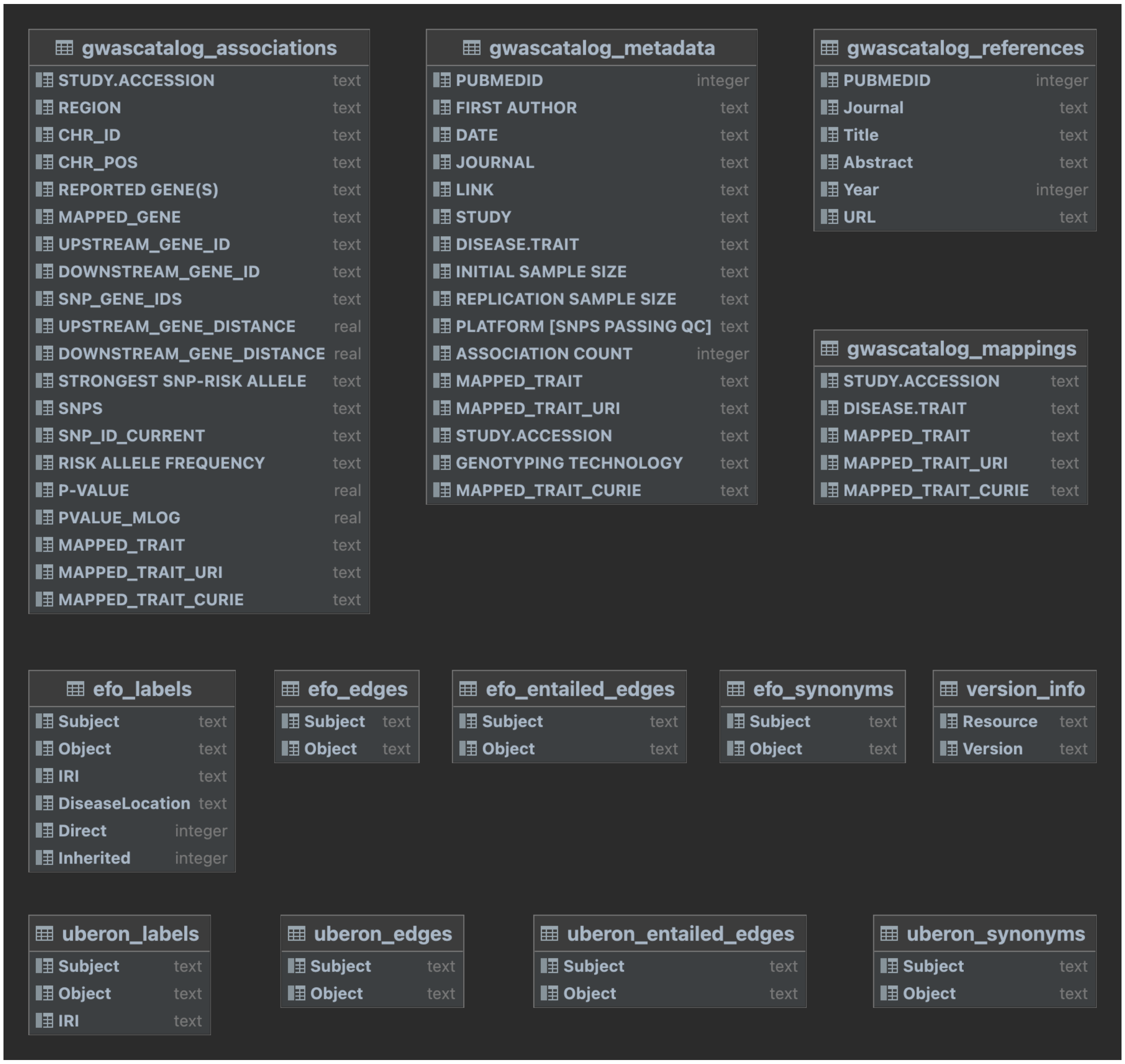


Figure 1: Table descriptions for GWAS catalog processed data.

#### Additional References

Xie, Yihui. 2024. *Bookdown: Authoring Books and Technical Documents with R Markdown*. Manual.
